# Distinguishing Critical, Beneficial, Neutral and Harmful Mutations Uncovered in the Directed Evolution of a Yeast Membrane Receptor

**DOI:** 10.1101/2020.08.04.236505

**Authors:** Adebola Adeniran, Keith E.J. Tyo

## Abstract

We present a reversion analysis of mutations introduced during the directed evolution of the yeast G-protein coupled receptor (GPCR) Ste2p to detect a peptide biomarker of chronic kidney disease. Two mutated receptors are analyzed in this study. Mutations acquired during directed evolution were reverted one at a time to the wild-type residue to assess the mutation’s contribution to receptor function. Mutations in the first and fifth transmembrane regions, the second intracellular loop and a truncation were found to be crucial for sensitive detection of the peptide biomarker. Some mutations acquired during directed evolution were found to be neutral to or harmful for biomarker detection. Mutations were also assessed for their contributions to increasing basal activity of the evolved receptors. A similar set of crucial mutations were found in the two receptors, implying a similar mechanism detection. The mutations are reasoned to appear to give the ability to detect a smaller sized peptide, affect interaction with the G-protein and allow for prolonged signaling after stimulation. These data should provide guidance for further engineering of Ste2p and other GPCRs.

## Introduction

Biosensing allows organisms to assess environmental conditions to make critical decisions concerning growth, reproduction and avoidance of harm through specific detection mechanisms. Biosensors take advantage of this precise and powerful biological specificity to detect and report the presence or absence of biological analytes of human interest such as biowarfare agents [1], [2], biofuel precursors [3], food contaminants [4], and environmental pathogens [5] among many others [6] .

However, creating biosensors for molecules for which there is no native sensing system to exploit remains challenging. In these cases, directed evolution can be used to generate a novel detection element for the analyte of interest. With the power to create novel detection elements, the possibilities for biosensors are endless. Though an exciting tool, directed evolution can yield mutations that are neutral or even harmful to the evolved function.

We previously reported a study on the directed evolution of the yeast GPCR Ste2p, which natively detects the α-factor peptide (amino acid sequence WHWLQLKPGQPMY), to detect an amidated peptide fragment of cystatin C (amino acid sequence ALDFAVGEYNK), as cystatin C is a biomarker for chronic kidney failure [7]. Directed evolution was a critical tool as no complete crystal structure of Ste2p has been solved to date that could be used to guide receptor engineering. The mutant receptors obtained from directed evolution contained many mutations away from the putative ligand binding site and it remained unclear which mutations played a role in the gain in function. Distinguishing which mutations play which roles can generate a working hypothesis as to how the mutant receptor is able to detect the cystatin peptide.

For two of the STE2 mutants produced from directed evolution, receptors Mut1 and Mut2, each mutation was individually reverted to the wild-type amino acid to quantify the effect of each individual mutation on 1) the receptor’s ability to detect the cystatin peptide and 2) the receptor’s increased basal activity. To quantify the role of each mutation on the ability to detect the cystatin peptide, the reversion-containing receptors were compared to the parent mutant receptor where all receptors were treated with the minimal amount of peptide needed to induce a response. To quantify the role of each mutation on the increase in basal activity, the reversion containing receptors were compared to the parent receptor where all receptors were untreated.

## Results

### Reversion of mutations

Receptor Mut1 contains 9 mutations as compared to WT Ste2p. The reversion containing receptors are labeled by the single reversion that distinguishes the receptor from Mut1. For example, Mut1-F26Y contains all the mutations in Mut1 except for mutation Y26F, which is reverted to the wildtype amino acid Y26. The only exception is receptor Mut1-Y158F. During the course of evolution, the N158Y mutation was mutated to and eventually mutated back to N158Y in the final Mut1 receptor. The Mut1-Y158F receptor was created to investigate the effect of the lost N158F mutation.

Receptor Mut2 contains 7 mutations as compared to WT Ste2p. The reversion containing receptors are labeled as described above. The notable exception is Mut2-del319F/325X. When the deletion and consequential truncation were reverted, two additional mutations after the truncation, Q328R and S356T, were allowed to be expressed. Because it was later determined that the receptor cannot detect the cystatin peptide without the deletion and truncation, Q328R and S356T were not further investigated. All receptor names and sequences are provided in Table 1.

**Table 1:** Receptor Sequences.

| Receptor | Mutations from WT Ste2p |
| --- | --- |
| Mut1 | Y26F,M54I,F55L,A61A*,R74K,N158Y,M218T,K225T,Y320X |
| Mut1-F26Y | M54I,F55L,A61A*,R74K,N158Y,M218T,K225T,Y320X |
| Mut1-I54M | Y26F,F55L,A61A*,R74K,N158Y,M218T,K225T,Y320X |
| Mut1-L55F | Y26F,M54I,A61A*,R74K,N158Y,M218T,K225T,Y320X |
| Mut1-A61A* | Y26F,M54I,F55L,R74K,N158Y,M218T,K225T,Y320X |
| Mut1-K74R | Y26F,M54I,F55L,A61A*,N158Y,M218T,K225T,Y320X |
| Mut1-Y158N | Y26F,M54I,F55L,A61A*,R74K,M218T,K225T,Y320X |
| Mut1-Y158F | Y26F,M54I,F55L,A61A*,R74K,N158F,M218T,K225T,Y320X |
| Mut1-T218M | Y26F,M54I,F55L,A61A*,R74K,N158Y,K225T,Y320X |
| Mut1-T225K | Y26F,M54I,F55L,A61A*,R74K,N158Y,M218T,Y320X |
| Mut1-X320Y | Y26F,M54I,F55L,A61A*,R74K,N158Y,M218T,K225T |
| Mut2 | D3N,M54I,N158Y,M218T,K225T,L228F,F319del/325X** |
| Mut2-N3D | M54I,N158Y,M218T,K225T,L228F,F319del/325X** |
| Mut2-I54M | D3N,N158Y,M218T,K225T,L228F,F319del/325X** |
| Mut2-Y158N | D3N,M54I,M218T,K225T,L228F,F319del/325X** |
| Mut2-T218M | D3N,M54I,N158Y,K225T,L228F,F319del/325X** |
| Mut2-T225K | D3N,M54I,N158Y,M218T,L228F,F319del/325X** |
| Mut2-F228L | D3N,M54I,N158Y,M218T,K225T,F319del/325X** |
| Mut2-del319F** | D3N,M54I,N158Y,M218T,K225T,L228F, (Q328R, S356T) *** |

\*A61A is a codon switch. \*\*The deletion at F319 caused an early truncation at amino acid 325. \*\*\*See text for details

First, critical mutations were identified. For receptor Mut1, the minimal amount of peptide to induce a response is 100 μM. Therefore, for receptor Mut1, critical mutations are defined as mutations that when reverted to the WT residue, there is no response to 100 μM ligand (i.e. there is no statistical difference between untreated and treated receptor containing the reversion of the critical mutation in the fluorescence-based assay described in Materials and Methods). Critical mutations in the Mut1 receptor are M54I, N158Y, M218T and Y320X as when these were removed, there was no response to ligand (Fig 1A.).

**Figure 1.**
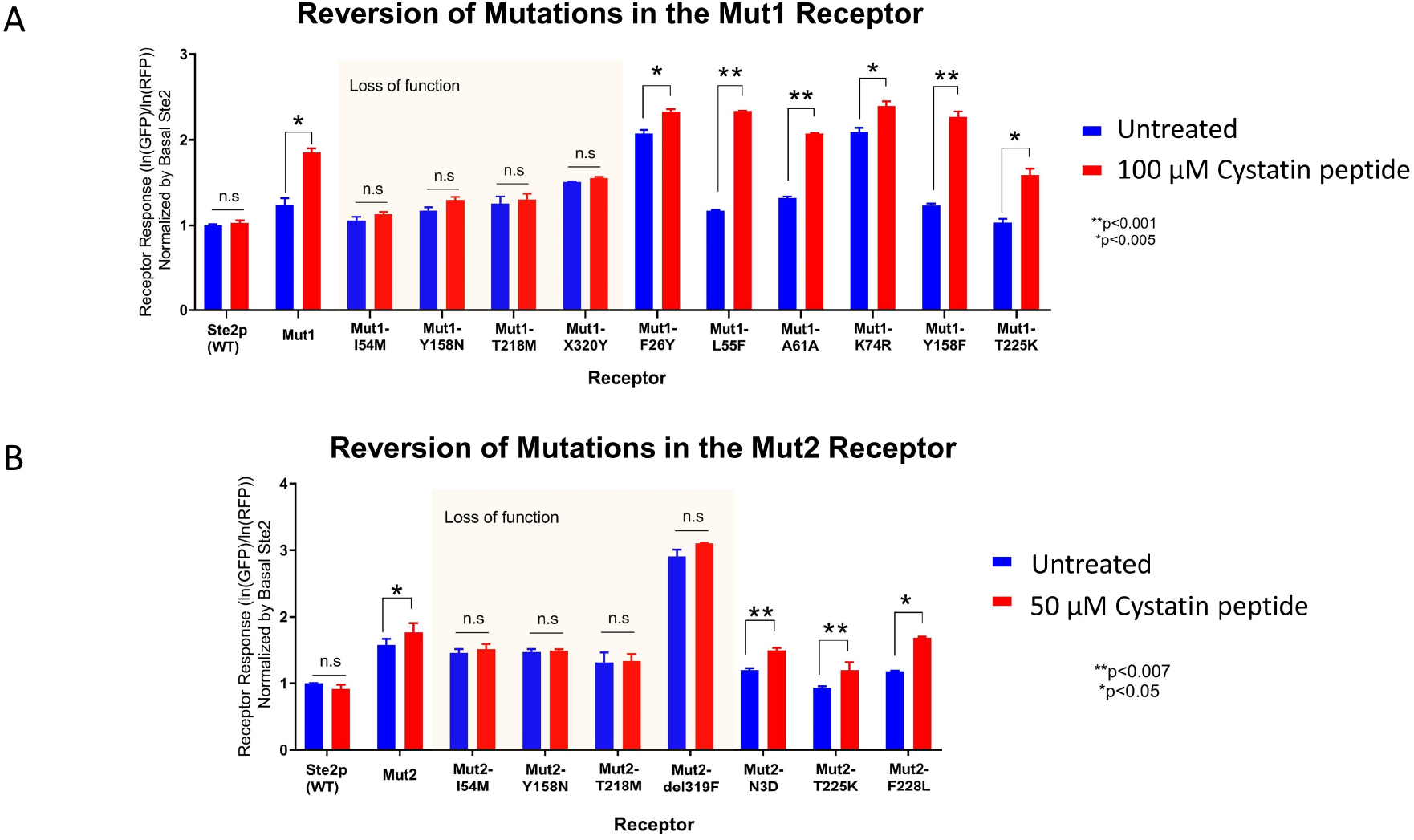
Reversion-containing Mutants Response to Cystatin Peptide. -**A**-Receptor Mut1 contains 9 mutations as compared to WT Ste2p. **B**-Receptor Mut2 contains 7 mutations as compared to WT Ste2p. The reversion containing receptors are labeled by the single reversion that distinguishes the receptor from the parent mutant receptor. See text for details. All receptor names and sequences are provided in Table 1. p-values are reported as determined by two-way ANOVA. Each data point is the average of 3 technical replicates. The experiment was replicated twice with comparable results.

For receptor Mut2, the minimal amount of peptide to induce a response is 50 μM. Therefore, for receptor Mut2, critical mutations are defined as mutations that when reverted, there is no response to 50 μM ligand. Critical mutations in the Mut2 receptor are M54I, N158Y, and M218T as when these were removed, there was no response to ligand (Fig 1B). The p-values for all critical mutations ranged from 0.08-0.99 as determined by 2 way ANOVA.

#### Peptide detection

Once critical mutations were identified, the remaining mutations were then classified as either harmful, beneficial or neutral. Harmful mutations meet two criteria: 1.) the receptor must still be able to respond to the cystatin ligand without the mutation and 2.) when the mutation is present, it reduces the receptor’s response to ligand. Beneficial mutations are not necessarily critical mutations (see above for definition of critical mutation), but do improve receptor response to ligand (i.e. the receptor becomes less sensitive to ligand when beneficial mutations are reverted to the WT residue). Neutral mutations neither improve nor hurt receptor sensitivity to ligand and there is no difference in receptor performance when the mutation is reverted to the WT residue.

Quantitatively, the classification of each mutation was determined by calculating two key parameters: the response of each reversion-containing mutant relative to that of the parent receptor (either Mut1 or Mut2) and the p-value of the responses. The relative responses and corresponding p-values are plotted against each other in Fig 2.

**Figure 2.**
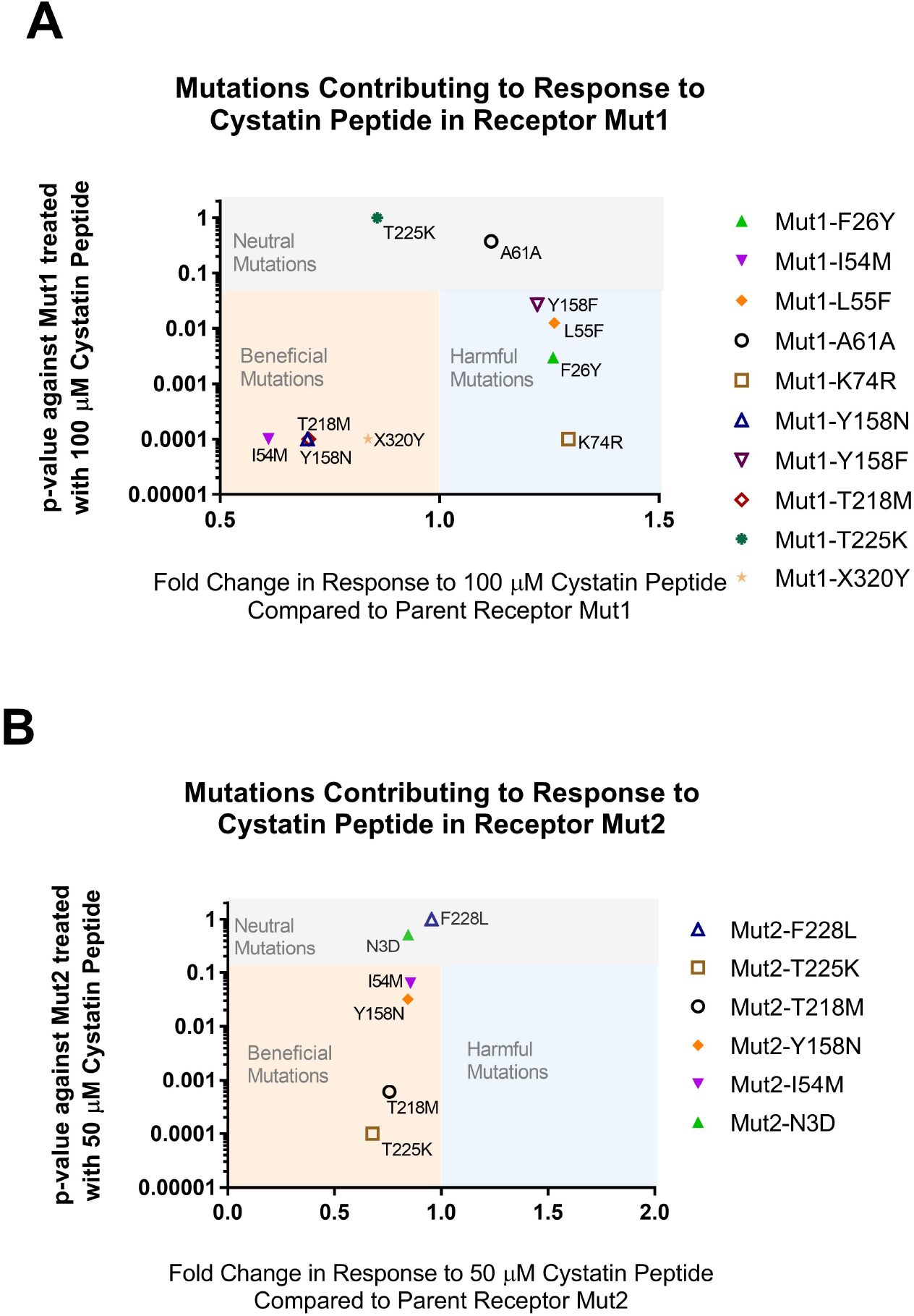
Mutations Contributing to Response to Cystatin Peptide in Mutant Receptors Mut1 (A) and Mut2 (B). The classification of each mutation was determined by calculating the response of each reversion-containing mutant relative to that of parent mutant receptor and plotting that against the statistical significance of the relative response. All receptor names and sequences are provided in Table 1. Each data point is the average of 3 technical replicates. The experiment was replicated twice with comparable results.

The classification of mutations was determined by clustering in the tornado plots in Fig 2. For the Mut1 data set, neutral mutations clustered at p-value> =0.100. All beneficial and harmful mutations had p-values <= 0.027. Specifically beneficial mutations gained are as: M54I, N158Y, M218T and Y320X. Harmful mutations are: Y26F, F55L, R74K and F158Y. (It should be noted that F158Y is labeled relative to an intermediate mutant that contained the N158F mutation. Mutation at position 158 from F to Y dampened response to peptide). Neutral mutations are K225T and the codon change at A61A.

For the Mut2 data set, neutral mutations clustered at p-value>= 0.100. All beneficial and harmful mutations clustered at p-<= 0.063. Specifically, beneficial mutations gained are identified as: M54I, N158Y, M218T, K225T and F319del (which resulted in a truncation at residue 325). There were no harmful mutations identified. Neutral mutations are D3N and F228L. A summary of all mutations is provided in Table 2.

**Table 2:** Roles of Mutations* in Affecting Response to Cystatin Peptide and Basal Activity.

|  | **Response to Cystatin Peptide** |  |  | **Basal Activity** |  |  |
| --- | --- | --- | --- | --- | --- | --- |
|  | Beneficial Mutations | Harmful Mutations | Neutral Mutations | Beneficial Mutations | Harmful Mutations | Neutral Mutations |
| Receptor Mut1 | **M54I**<br>**N158Y**<br>**M218T**<br>**Y320X** | Y26F<br>F55L<br>R74K<br>F158Y | A61A<br>K225T | M54I<br>N158Y<br>K225T | Y26F<br>R74K<br>Y320X | F55L<br>A61A<br>F158Y<br>M218T |
| Receptor Mut2 | **M54I**<br>**N158Y**<br>**M218T**<br>K225T<br>**F319del-325X** | none | D3N<br>L228F | M218T<br>K225T | F319del-325X | D3N<br>M54I<br>N158Y<br>L228F |

\*Mutations critical for response to the cystatin peptide are in bold.

#### Basal activity

Over the course of directed evolution, the basal activity of both mutant receptors increased modestly. Many individual mutations appeared to have an effect on basal activity. The increase in basal activity was interrogated to identify causative mutations. As before, mutations were classified as either harmful, beneficial or neutral towards basal activity. For the Mut1 receptor, beneficial mutations for basal activity are M54I, K225T and N158Y. Harmful mutations for basal activity are F26Y, R74K and Y320X. Neutral mutations for basal activity are L55F, A61A, M218T and F158Y (Fig 3A). For the Mut2 receptor, beneficial mutations for basal activity are M218T and K225T. The only harmful mutation for basal activity is a deletion at F319 that resulted in a premature truncation at position 325. Neutral mutations for basal activity are D3N, M54I, N158Y and F228L (Fig 3B). A summary of all mutations is provided in Table 2.

**Figure 3.**
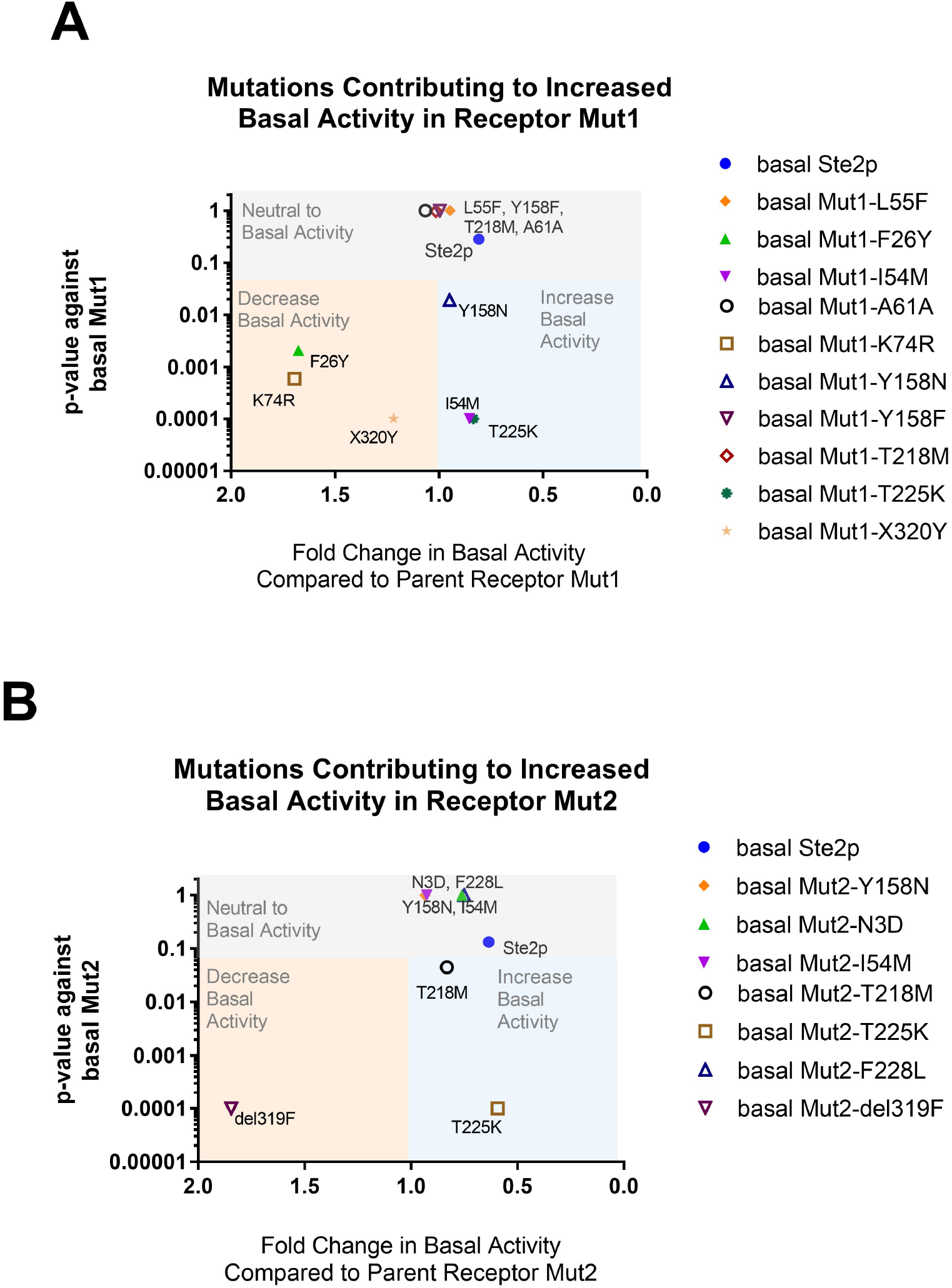
Mutations Contributing to Basal Activity in Mutant Receptors. The classification of each mutation was determined by calculating the basal activity of each reversion containing mutant relative to that of receptor Mut1 (**A**) or Mut2 (**B**)and comparing the relative basal activity to the statistical significance of the relative activity. All receptor names and sequences are provided in Table 1. Each data point is the average of 3 technical replicates. The experiment was replicated twice with comparable results.

## Discussion

### Peptide detection

#### Critical and beneficial mutations for peptide detection

Though information on sequence-structure relationships is hard to come by for many GPCRs, Ste2p is unique as there is a rich literature on this topic. Here, the mutations in receptors Mut1 and Mut2 are considered in context of this rich literature. A visualization of all mutations is provided in the snake plot in Fig 4. Mutations M54I, N158Y and M218T were both critical mutations in receptors Mut1 and Mut2. Additionally, truncations at residues 320 and 325 were critical to both receptors. Mutation M54I was gained during the first round of evolution to a chimeric peptide similar to α-factor (Cys1) (S1 Table), and has been reported to give Ste2p the ability to respond to the synthetic α-factor variant desTrp^1^[Ala^3^,Nle^12^]α-factor [8] which is similar in size and sequence to the chimeric Cys1 ligand.

**Figure 4.**
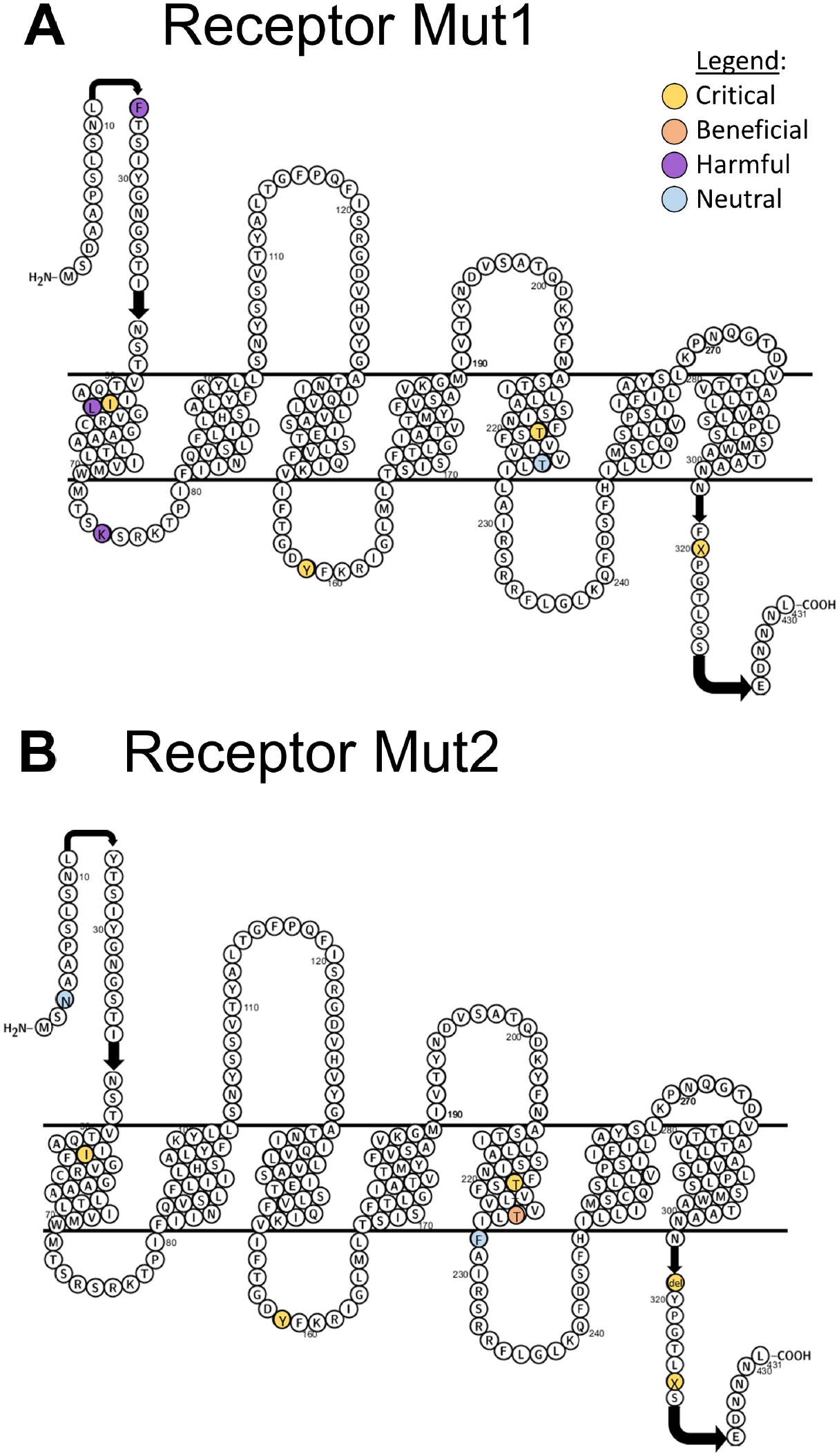
Snake plot diagram showing locations of mutations acquired during directed evolution. No mutations were evolved in extracellular loop 2, or transmembrane regions 2,3 6 or 7. Mutations are color-coded by whether they are critical, beneficial, harmful or neutral for detection of the cystatin peptide. Snake plot generated with Protter [9].

Amino acid 158 is mutated throughout the directed evolution pathway from an amide residue (N) to increasingly hydrophobic residues (Y and F). Previous work has indicated hydrophobic changes in the second intracellular loop may assist with G-protein coupling as the second intracellular loop is accessible to the G-protein [10].

M218T is located in the fifth transmembrane region, the relative motion of which is required for receptor activation [11]. In the context of the native Ste2p, M218T on its own is a neutral mutation, and residue 218 is believed to interact with residues on the third transmembrane region. Notably, M218T can rescue receptor functionality when paired with function-ablating mutations in the third transmembrane region [12], and a mutation in this position may help the receptor cope with otherwise destabilizing mutations.

Truncations on the C-terminal tail were critical for both receptors. Truncation Y320X is critical to receptor Mut1, and a deletion at residue 319 resulting in a truncation at residue 325 is critical to receptor Mut2. The Y320X truncation in receptor Mut1 was gained during evolution for peptide Cys4 (a chimera sharing intermediate sequence similarity to both α-factor and the cystatin peptide), and the 325X truncation in receptor Mut2 was gained during evolution for peptide Cys5, which differs from peptide Cys4 by two amino acid residues. The GPCR tail, where these mutations lie, contains multiple ubiquitination sites necessary for receptor endocytosis [13] and signal desensitization [14] following ligand stimulation. Thus, these truncations are predicted to provide prolonged signaling after ligand stimulation.

With the exception of the premature truncation at residue 325, all critical mutations in receptors Mut1 and Mut2 appeared during evolution for peptides which differ from the native α-factor in the signaling region of the peptide (closer to the N terminus of the peptide), but not in the binding region of the peptide. This may contribute to the fact that receptors Mut1 and Mut2 cannot respond to peptides that have different binding regions than α-factor unless the ligand is amidated on the C terminus.

The Mut2 receptor benefited from the K225T mutation, but the mutation was not critical for response to the cystatin peptide. In receptor Mut1, the K225T mutation was neutral regarding peptide response. In another study, the K225C mutation ablated functionality of the native Ste2p receptor [15], so it is noteworthy that a substitution to a similarly nucleophilic residue (K225T) did not ablate receptor functionality here. K225T is on the border of the third intracellular loop and the fifth transmembrane region, the relative motion of which is required for receptor activation [11]. If the K225T mutation is destabilizing, a possible explanation for the beneficial nature of the K225T mutation in the Mut2 receptor is that the mutation destabilizes the receptor enough to accept a non-native ligand, but the M218T mutation, which was noted as a rescuing mutation earlier in this discussion, rescues this destabilization.

Interestingly, the K225T mutation is neutral in receptor Mut1. Given the similarities of the other critical mutations between both receptors, it is surprising that K225T is neutral in Mut1 but beneficial in Mut2. The difference in critical mutations between the two receptors is in the truncations; however, both truncations result in the loss of the same number of predicted ubiquitination sites [16]. As K225T and the C-terminal tail face the intracellular region, they could potentially interact with downstream effectors. More biophysical studies examining interactions between network proteins and GPCR structure would need to be performed before drawing firm conclusions on the differences of mutation K225T in receptors Mut1 and Mut2.

In summary, the three critical mutations common to both receptors, M54I, N158Y, and M218T appear to give the ability to detect a smaller sized peptide, alter interaction with the G-protein and allow for prolonged signaling after stimulation, respectively. Our observations are consistent with previous findings on GPCR mechanism. In the context of the native Ste2p receptor, residues 156-162 can be removed without drastically affecting response to the native α-factor [17]. However, the findings of this work support both the formation of a hydrophobic pocket by intracellular loops to interact with the G-protein [10], and the importance of the intracellular loops in mediating signaling from C-terminal truncated receptors [18]. Because the C-terminus interacts with the G-protein [19], C-terminal truncated receptors may become more dependent on intracellular loop contacts with the G-protein though this remains to be determined. Residue 158 is accessible to the G-protein [10], and the role of the second intracellular loop in coupling to G-proteins in other GPCR families is documented [20]–[22] . Finding beneficial mutations far from the putative binding site is not surprising as it has been demonstrated that Ste2p can be evolved to detect non-native peptides through the alteration of interactions with other proteins in the signaling network [23].

#### Harmful mutations for peptide detection

For the Mut1 receptor, mutations Y26F, F55L, R74K and F158Y were found to be detrimental for detection of the cystatin peptide. There were no detrimental mutations found for the Mut2 receptor. It is noteworthy that Y26F, R74K, and F158Y, all harmful mutations for cystatin detection, appeared in the same round of directed evolution, but also co-occurred with critical mutation Y320X. The mutations believed to be important for global stability, M218T and K225T, had appeared in previous generations, which may have allowed for the accommodation of the harmful mutations.

Oligomerization has been reported to play a role in the formation of ligand binding sites [24]–[26] and signal transduction [27], [28] in a variety of GPCRs. The Y26F mutation is located in the N terminus, which has been reported to undergo a conformational change upon ligand binding and be involved in receptor dimerization in Ste2p. Residue 26, along with residues 22 and 28, form the dimer interface of the N terminus [29]. A Y26C Ste2p mutant was reported to have an 80% decrease in surface expression levels [29]. If any mutation at residue 26 is unfavorable, then the Y26F mutation may also cause decreased surface expression, reducing the number of receptors available to detect peptide and initiate downstream signaling. The Y26F mutation also decreased basal signaling activity, indicating that this position influences receptor signaling.

The F55L mutation is harmful for the response of receptor Mut1 to the cystatin peptide. Residue 55 has been implicated in ligand specificity and signal transduction [30]. Specifically, the F55V mutation causes decreased signaling and reduces binding affinity to the native α-factor by over 10-fold. Given the hydrophobic nature of both valine and leucine, it is possible that the F55L mutation has a similar effect as the F55V mutation and causes decreased signaling. Also noteable is the first transmembrane region plays a key role in receptor dimerization, although residue 55 is predicted to face away from the dimerization interface [31].

The R74K mutation is harmful for the response of receptor Mut1 to the cystatin peptide and is located in the first intracellular loop. The R74S and R76S mutations cause decreased signaling in response to α-factor for Ste2p [32]. This could implicate the importance of strongly charged residues in the first intracellular loop. Although lysine is a basic residue, its pKa (10.79) is lower than that of arginine (12.48), thus the strength of the basic charge is likely important at residue 74. To our knowledge, a specific role of the first intracellular loop has not been explicitly defined for Ste2p, however, the first intracellular loop is involved in transport from ER to cell surface in the α_2B_-adrenergic receptor [33]. A mutation in this region may disrupt receptor trafficking in Ste2p, though more experiments would need to be done to confirm this hypothesis. The R74K mutation is also harmful for basal activity, which indicates that this residue may play a role in signal transduction.

The importance of position 158 was discussed in the previous section “Critical and Beneficial Mutations for Peptide Detection”. The native Ste2p has N at position 158. Over the course of evolution, N158 was first mutated to Y, then to F, then again to Y. That the F158Y mutation was harmful indicates that F is preferred at this position to Y with respect to detecting the cystatin peptide.

#### Neutral mutations for peptide detection

For receptor Mut1, mutations A61A (codon switch) and K225T were neutral with regards to detecting the cystatin peptide. It is not surprising that the codon change did not affect response to peptide. It is interesting, however, that the K225T mutation was neutral in the Mut1 receptor, but beneficial in the Mut2 receptor. For receptor Mut2, mutations D3N and F228L were neutral with regards to detecting the cystatin peptide. The similar hydrophobic nature of F and L may explain the neutral effect of the F228L substitution. To our knowledge, there is no precedent in literature for mutations at D3 affecting receptor response to ligand.

### Basal activity

#### Beneficial mutations for increased basal activity

We define basal activity as ligand-independent signaling. Mutations that affect basal activity are implicated in receptor signaling. For receptor Mut1, mutations M54I, N158Y and K225T increased basal activity. Interestingly, mutations M54I and N158Y were also crucial for response to the cystatin peptide. For receptor Mut2, mutations M218T and K225T increased basal activity and were also both beneficial for detecting the cystatin peptide. While a generalized increase in basal activity has not been explicitly shown to facilitate detection of non-native ligands for Ste2p, an overlap in mutations that increase basal activity and are also beneficial for peptide detection suggests a mechanism where a generalized increase in basal activity facilitates detection of non-native ligands. This is a feasible hypothesis if mutations that increase basal activity simultaneously increase promiscuity. Promiscuity is often observed during directed evolution and becomes a starting point for evolving a new protein function [34]. If the mutations that increase basal activity also increase promiscuity, then a generalized increase in basal activity may facilitate detection of non-native ligands. Additionally, specificity can be altered through the interaction of the receptor with downstream effectors [23]. The overlap between mutations that increase basal activity and those that are crucial and beneficial for detection of the cystatin peptide suggests that an increase in basal activity was necessary along the evolutionary pathway taken in this directed evolution study.

#### Harmful mutations for basal activity

For receptor Mut1, the Y320X mutation is critical for peptide response but detrimental to increasing basal activity. Mutations Y26F and R74K were detrimental to both peptide detection and basal activity, suggesting that mutations at these sites could affect overall receptor signaling. Mutations Y26F and R74K were discussed in the previous section “Harmful Mutations for Peptide Detection”. For receptor Mut2, the F319del/325X mutation was harmful for both detection of the cystatin peptide and basal activity. In both receptors, premature truncations were harmful for basal activity. This implies that the presence of the truncation reduces basal activity. More biophysical studies would need to be carried out to determine how the truncations impact signaling effectors.

#### Neutral mutations for basal activity

For receptor Mut1, mutations L55F, A61A, F158Y and M218T were neutral for basal activity. The M218T mutation was beneficial, but not crucial, for response to peptide. That the M218T mutation does not affect basal activity implies that the mutation is specific for increasing sensitivity towards the cystatin peptide. For receptor Mut2, mutations D3N, M54I, N158Y and F228L were neutral for basal activity. D3N and F228L were also neutral with respect to peptide response. That mutation N158Y was critical for response of Mut2 to the cystatin peptide but is neutral with regards to basal activity implies that the N158Y mutation specifically affects receptor specificity in the Mut2 receptor. This is in contrast to the Mut1 receptor, where the N158Y mutation was crucial for response to peptide and increasing basal activity.

### Additional comments on mutations

It is noteworthy that the Q272Y mutation was lost over the course of directed evolution. This mutation is located in the third extracellular loop, which is important for receptor specificity [35]. Interestingly, no mutations were gained in the second or fourth transmembrane regions. It is believed that there are common underlying mechanisms between class A GPCRs, typically represented by the rhodopsin receptor, and class D GPCRs, of which Ste2p is a member [36]. The second or fourth transmembrane regions, which are believed not to change structurally during receptor activation in class A GPCRs [36], may provide similar structural support in class D GPCRs. Additionally, the two receptor classes share highly conserved microdomains on the sixth and seventh transmembrane regions. That we did not observe any mutations in these transmembrane regions may be explained by the conservation of these regions. We were surprised that we did not observe mutations in the second extracellular loop as this region has been implicated in signaling and specificity [37].

## Conclusions

Overall, the new receptor function is a result of mutations that do not solely affect ligand binding. Over the course of directed evolution, a mutation near the binding pocket (M54I) was observed first, followed by mutations that are known to compensate for detrimental mutations (M218T, K225T), and mutations that may affect interaction with other signaling network proteins (truncations, N158Y). It has also been demonstrated that the response of Ste2p and mutant receptors to non-native peptides can be regulated through altered interactions in the pathway signaling network, specifically in C-terminally truncated receptors. The C-terminal truncations supposedly lowered the energetic threshold for activation by disrupting the interaction of the receptor and the downstream pathway effectors such as Sst2 [23]. C-terminal truncations were crucial for detecting the cystatin peptide in this work. Future experiments could be carried out to clarify what role the C-terminal truncation plays in the detection of the cystatin peptide. Sst2 is knocked out in the strain used for biosensor development, so the novel detection capabilities could not have been gained through an altered interaction with Sst2. However, there are many other pathway effectors, including the G protein the Ste5/Ste11 and Ste20 complexes [38], that could be investigated for altered interactions.

In summary, we found mutations that were critical to the mutant receptors function, some that were beneficial, but not critical to the mutant receptor function, and some that were neutral or deleterious to the mutant receptors function. The mutations were largely consistent with known literature on sequence-function relationships for Ste2p and could guide more focused receptor engineering efforts.

## Materials and Methods

### Strains and media

Yeast strain MPY578t5 was a gift from Bryan Roth. STE2 was obtained from the genomic DNA of strain ESM356-1 and yeGFP and natNT2 were obtained from plasmid pCT191, which were both kindly given from Michael Knop’s lab. p416GPD from the Mumberg plasmids was obtained from Addgene. All oligonucleotides were synthesized by Integrated DNA Technologies (Coralville, USA). Alpha-factor mating pheromone was purchased from Sigma-Aldrich (St. Louis, USA). All C-terminally amidated peptides were kindly synthesized by the Mrksich lab at Northwestern University (Evanston, USA).

Yeast with the uracil auxotrophy were cultivated in complete synthetic media without uracil prepared with -His-Ura dropout supplement (Clontech, Mountain View, CA) according to manufacturer’s instructions and supplemented with 20 g/mL histidine and 80mg/L adenine hemisulfate (Sigma-Aldrich, St. Louis, USA)

### Cloning

Cloning of strains was previously reported in [7]. Briefly, strains contain yeGFP at the FUS1 locus to allow for fluorescent reporting of receptor activation and constitutively express mKate. Specifically, the reversions were examined in background strain yJB013. Transformations were done via a standard chemical method [39].

The NEBaseChanger online tool (available at https://nebasechanger.neb.com/) was used to design primers to revert mutations to the wild-type sequence at that location. The primers were used to linearize plasmids containing the mutant yeast sequences. The linearized plasmids were cleaned with the Zymo Clean and Concentrate kit and eluted in 30 μL of nuclease-free water. The cleaned DNA was then used in the KLD Enzyme Kit (New England Biolabs) with 1 μL KLD Enzyme Mix, 5 μL 2× KLD Buffer, and 4 μL cleaned PCR product. The mix incubated at room temperature for 10-30 minutes before being transformed into chemically competent DH5α cells. Plasmids were isolated and sequenced verified before being transformed into yeast. Receptors were again sequence verified by colony PCR after transformation into yeast.

### Assays

Yeast were grown overnight at 30 °C and 225 rpm in 5 mL culture tubes and then diluted to an OD600 = 0.1 in CSM-Ura media . Immediately after dilution, the peptide ligand was added. The culture was incubated for 2.5 h at 30°C with 225 rpm shaking. Cells were centrifuged at 2000g for 5 min and supernatant was removed. Cells were resuspended in 1× PBS in preparation for flow cytometry. Analytical flow cytometry was performed with the BD FORTRESSA (BD Biosciences, San Jose, USA); yEGFP was read on the FITC-channel, and mKate was read on the PE-Texas Red channel. Receptor output was measured by dividing the geometric mean of GFP fluorescence by the geometric mean of RFP fluorescence for a population. Specifically, basal activity is measured as ligand-independent signaling.

## Acknowledgements

This work is funded by National Science Foundation Grant DGE-1324585, the Bill and Melinda Gates Foundation Grant OPP1061177, the Northwestern University Presidential Fellowship, and the Howard Hughes Medical Institute Gilliam Fellowship for Advanced Study. We thank Bryan Roth at the University of North Carolina-Chapel Hill for the gift of the MPY578t5 yeast strain, Alexei Tan at Northwestern University for help in synthesizing peptides, the NU Seq Core (Northwestern University), and the Northwestern University Robert H. Lurie Flow Cytometry Core Facility.

## Supporting information

S1 Table. **Chimeric ligands used to traverse the evolutionary landscape in a step-wise fashion.** See [7] for details. The sequences of the native ligand, α factor, and chimeric ligands are provided.

S2 Table. **Statistical significances for Mut1 reversion experiments**. P-values for experiments with parent receptor Mut1 are provided as determined by 2 way ANOVA.

S3 Table. **Statistical significances for Mut2 reversion experiments.** P-values for experiments with parent receptor Mut2 are provided as determined by 2-way ANOVA.

## References

[1] S. S. Iqbal, M. W. Mayo, J. G. Bruno, B. V. Bronk, C. A. Batt, and J. P. Chambers, “A review of molecular recognition technologies for detection of biological threat agents,” Biosens. Bioelectron., vol. 15, no. 11–12, pp. 549–578, 2000.

[2] V. Radhika, T. Proikas-Cezanne, M. Jayaraman, D. Onesime, J. H. Ha, and D. N. Dhanasekaran, “Chemical sensing of DNT by engineered olfactory yeast strain,” Nat. Chem. Biol., vol. 3, no. 6, pp. 325–330, 2007.

[3] K. Mukherjee, S. Bhattacharyya, and P. Peralta-Yahya, “GPCR-Based Chemical Biosensors for Medium-Chain Fatty Acids,” ACS Synth. Biol., vol. 4, no. 12, pp. 1261–1269, 2015.

[4] M. Gonchar, O. Smutok, M. Karkovska, N. Satsyuk, and G. Gayda, “Yeast-Based Biosensors for Clinical Diagnostics and Food Control,” in Biotechnology of Yeasts and Filamentous Fungi, p. 391.

[5] N. Ostrov et al., “A modular yeast biosensor for low-cost point-of-care pathogen detection,” Sci. Adv., vol. 3, no. 6, p. e1603221, 2017.

[6] A. Adeniran, M. Sherer, and K. E. J. Tyo, “Yeast-based biosensors: Design and applications,” FEMS Yeast Res., vol. 15, no. 1, pp. 1–15, 2015.

[7] A. Adeniran, S. Stainbrook, J. W. Bostick, and K. E. J. Tyo, “Detection of a Peptide Biomarker by Engineered Yeast Receptors,” ACS Synth. Biol., vol. 7, no. 2, pp. 696–705, 2018.

[8] L. Marsh, “Substitutions in the Hydrophobic Core of the a-Factor Receptor of Saccharomyces cerevisiae Permit Response to Saccharomyces kluyveri ac-Factor and to Antagonist,” Mol. Celluar Biol., vol. 12, no. 9, pp. 3959–3966, 1992.

[9] U. Omasits, C. H. Ahrens, S. Müller, and B. Wollscheid, “Protter: Interactive protein feature visualization and integration with experimental proteomic data,” Bioinformatics, vol. 30, no. 6, pp. 884–886, 2014.

[10] Y. Choi, J. B. Konopka, and S. Brook, “Accessibility of Cys residues substituted into the cytoplasmic regions of the α-factor receptor defines residues that are accessible for G protein activation †,” Biochemistry, vol. 45, pp. 15310–15317, 2006.

[11] A. Taslimi, E. Mathew, A. Ćelić, S. Wessel, and M. E. Dumont, “Identifying functionally important conformational changes in proteins: Activation of the yeast α-factor receptor Ste2p,” J. Mol. Biol., vol. 418, no. 5, pp. 367–378, 2012.

[12] C. M. Sommers and M. E. Dumont, “Genetic interactions among the transmembrane segments of the G protein coupled receptor encoded by the yeast STE2 gene.,” J. Mol. Biol., vol. 266, no. 3, pp. 559–575, 1997.

[13] L. Hicke and H. Riezman, “Ubiquitination of a yeast plasma membrane receptor signals its ligand-stimulated endocytosis.,” Cell, vol. 84, no. 2, pp. 277–87, Jan. 1996.

[14] E. Leberer, D. Y. Thomas, and M. Whiteway, “Pheromone signalling and polarized morphogenesis in yeast,” Curr. Opin. Genet. Dev., vol. 7, no. 1, pp. 59–66, 1997.

[15] P. Dube, a DeCostanzo, and J. B. Konopka, “Interaction between transmembrane domains five and six of the alpha-factor receptor.,” J. Biol. Chem., vol. 275, no. 34, pp. 26492–26499, 2000.

[16] P. Radivojac et al., “Identification, analysis, and prediction of protein ubiquitination sites,” Proteins Struct. Funct. Bioinforma., vol. 78, no. 2, pp. 365–380, 2010.

[17] N. P. Martin, L. M. Leavitt, C. M. Sommers, and M. E. Dumont, “Assembly of G protein-coupled receptors from fragments: Identification of functional receptors with discontinuities in each of the loops connecting transmembrane segments,” Biochemistry, vol. 38, no. 2, pp. 682–695, 1999.

[18] S. L. Chinault, M. C. Overton, and K. J. Blumer, “Subunits of a Yeast Oligomeric G Protein-coupled Receptor Are Activated Independently by Agonist but Function in Concert to Activate G Protein Heterotrimers,” J. Biol. Chem., vol. 279, no. 16, pp. 16091–16100, 2004.

[19] M. Dosil, K. a Schandel, E. Gupta, D. D. Jenness, and J. B. Konopka, “The C terminus of the Saccharomyces cerevisiae alpha-factor receptor contributes to the formation of preactivation complexes with its cognate G protein.,” Mol. Cell. Biol., vol. 20, no. 14, pp. 5321–5329, 2000.

[20] A. J. Venkatakrishnan, X. Deupi, G. Lebon, C. G. Tate, G. F. Schertler, and M. M. Babu, “Molecular signatures of G-protein-coupled receptors,” Nature, vol. 494, no. 7436, pp. 185–194, 2013.

[21] E. S. Burstein, T. a. Spalding, and M. R. Brann, “The second intracellular loop of the m5 muscarinic receptor is the switch which enables G-protein coupling,” J. Biol. Chem., vol. 273, no. 38, pp. 24322–24327, 1998.

[22] J. Gomeza, C. Joly, R. Kuhn, T. Knöpfel, J. Bockaert, and J. P. Pin, “The second intracellular loop of metabotropic glutamate receptor 1 cooperates with the other intracellular domains to control coupling to G-proteins,” J. Biol. Chem., vol. 271, no. 4, pp. 2199–2205, 1996.

[23] R. B. Di Roberto, B. Chang, A. Trusina, and S. G. Peisajovich, “Evolution of a G protein-coupled receptor response by mutations in regulatory network interactions,” Nat. Commun., vol. 7, p. 12344, 2016.

[24] H. E. H. and R. Z. L T Potter, L A Ballesteros, L H Bichajian, C A Ferrendelli, A Fisher, “Evidence of Paired M2 Muscarinic Receptors,” Pharmacology, vol. 39, no. 2, pp. 211–221, 1991.

[25] B. A. Jordan and L. A. Devi, “G-protein-coupled receptor heterodimerization modulates receptor function,” Nature, vol. 399, no. 6737, pp. 697–700, 1999.

[26] A. Mijares, D. Lebesgue, G. Wallukat, and J. Hoebeke, “From agonist to antagonist: Fab fragments of an agonist-like monoclonal anti-β2-adrenoceptor antibody behave as antagonists.,” Mol. Pharmacol., vol. 58, no. 2, pp. 373–9, 2000.

[27] T. E. Hebert et al., “A peptide derived from a beta2-adrenergic receptor transmembrane domain inhibits both receptor dimerization and activation.,” J. Biol. Chem., vol. 271, no. 27, pp. 16384–16392, 1996.

[28] S. AbdAlla, H. Lother, and U. Quitterer, “AT1-receptor heterodimers show enhanced G-protein activation and altered receptor sequestration,” Nature, vol. 407, no. 6800, pp. 94–98, 2000.

[29] M. S. Uddin, H. Kim, A. Deyo, F. Naider, and J. M. Becker, “Identification of residues involved in homodimer formation located within a β-strand region of the N-terminus of a Yeast G protein-coupled receptor,” J. Recept. Signal Transduct., vol. 32, no. 2, pp. 65–75, 2012.

[30] M. G. Abel, B. K. Lee, F. Naider, and J. M. Becker, “Mutations affecting ligand specificity of the G-protein-coupled receptor for the Saccharomyces cerevisiae tridecapeptide pheromone,” Biochim. Biophys. Acta - Mol. Cell Res., vol. 1448, no. 1, pp. 12–26, 1998.

[31] M. C. Overton, S. L. Chinault, and K. J. Blumer, “Oligomerization, biogenesis, and signaling is promoted by a glycophorin A-like dimerization motif in transmembrane domain 1 of a yeast G protein-coupled receptor.,” J. Biol. Chem., vol. 278, no. 49, pp. 49369–49377, 2003.

[32] A. Ćelić, N. P. Martin, C. D. Son, J. M. Becker, F. Naider, and M. E. Dumont, “Sequences in the intracellular loops of the yeast pheromone receptor ste2p required for G protein activation,” Biochemistry, vol. 42, no. 10, pp. 3004–3017, 2003.

[33] M. T. Duvernay, C. Dong, X. Zhang, M. Robitaille, T. E. Hébert, and G. Wu, “A single conserved leucine residue on the first intracellular loop regulates ER export of G protein-coupled receptors,” Traffic, vol. 10, no. 5, pp. 552–566, 2009.

[34] J. D. Bloom and F. H. Arnold, “In the light of directed evolution: Pathways of adaptive protein evolution,” Proc. Natl. Acad. Sci., vol. 106, no. Supplement_1, pp. 9995–10000, 2009.

[35] M. Sen, A. Shah, and L. Marsh, “Two types of alpha-factor receptor determinants for pheromone specificity in the mating-incompatible yeasts S. cerevisiae and S. kluyveri,” Curr. Genet., vol. 31, no. 3, pp. 235–240, 1997.

[36] M. Eilers, V. Hornak, S. O. Smith, and J. B. Konopka, “Comparison of class A and D G protein-coupled receptors: Common features in structure and activation,” Biochemistry, vol. 44, no. 25, pp. 8959–8975, 2005.

[37] J. C. Lin, W. Parrish, M. Eilers, S. O. Smith, and J. B. Konopka, “Aromatic residues at the extracellular ends of transmembrane domains 5 and 6 promote ligand activation of the G protein-coupled α-factor receptor,” Biochemistry, vol. 42, no. 2, pp. 293–301, 2003.

[38] L. Bardwell, “A walk-through of the yeast mating pheromone response pathway,” Peptides, vol. 26, no. 2, pp. 339–350, Feb. 2005.

[39] R. D. Gietz and R. H. Schiestl, “Large-scale high-efficiency yeast transformation using the LiAc/SS carrier DNA/PEG method.,” Nat. Protoc., vol. 2, no. 1, pp. 38–41, 2007.

