## Supplemental Tables for "Distinguishing Critical, Beneficial, Neutral and Harmful Mutations Uncovered in the Directed Evolution of a Yeast Membrane Receptor"

**Supplementary Table 1. Chimeric ligands used to traverse the evolutionary landscape in a step-wise fashion**. The sequences of the native ligand, α factor, and chimeric ligands are provided.

| Name | Sequence |
|---|---|
| α-factor | WHWLQLKPGQPMY |
| Cys1 | - HALALKPGEPMY |
| Cys2 | - - ALALKPGEPMY |
| Cys3 | - - ALDFKPGEPMY |
| Cys4 | - - ALDLAVGEPMY |
| Cys5 | - - ALDFAVGEPMK |
| Cys6 | - - ALDFAVGEYMK |
| Cys 7 – Cystatin peptide | - - ALDFAVGEYNK |

**Supplementary Table 2. Statistical significances for Mut1 reversion experiments.** P-values for experiments with parent receptor Mut1 are provided as determined by 2-way ANOVA.

| Receptor | p-value: receptor treated with 100 µM cystatin peptide vs receptor untreated | p-value: receptor treated with 100 µM cystatin peptide vs Mut1 treated with 100 µM cystatin peptide | p-value: receptor untreated vs Mut1 untreated |
|---|---|---|---|
| Ste2 | >0.9999, ns | 0.0009 | 0.2817 |
| Mut1 | <0.0001 | -- | -- |
| Mut1-F26Y | <0.0001 | 0.0030 | 0.0021 |
| Mut1-I54M | 0.8400, ns | <0.0001 | <0.0001 |
| Mut1-L55F | 0.0002 | 0.0126 | >0.9999 |
| Mut1-A61A | 0.0015 | 0.3759 | >0.9999 |
| Mut1-K74R | <0.0001 | 0.0001 | 0.0006 |
| Mut1-Y158N | 0.0842, ns | <0.0001 | 0.0192 |
| Mut1-Y158F | 0.0003 | 0.0270 | >0.9999 |
| Mut1-T218M | >0.9999, ns | <0.0001 | 0.9694 |
| Mut1-T225K | <0.0001 | >0.9999, ns | <0.0001 |
| Mut1-X320Y | 0.9870, ns | <0.0001 | <0.0001 |

**Supplementary Table 3. Statistical significances for Mut2 reversion experiments.** P-values for experiments with parent receptor Mut2 are provided as determined by 2-way ANOVA.

| Receptor | p-value: receptor treated with 50 µM cystatin peptide vs Mut2 treated with 50 µM cystatin peptide | p-value: receptor treated with 50 µM cystatin peptide vs receptor untreated | p-value: receptor untreated vs Mut2 untreated |
|---|---|---|---|
| Ste2 | 0.0007 | >0.9999, ns | 0.1335 |
| Mut2 | -- | 0.0062 | -- |
| Mut2-N3D | 0.5215, ns | 0.0348 | >0.9999, ns |
| Mut2-I54M | 0.0632 | >0.9999 | 0.9969 |
| Mut2-Y158N | 0.0321 | >0.9999 | >0.9999, ns |
| Mut2-T218M | 0.0006 | >0.9999 | 0.0448 |
| Mut2-T225K | <0.0001 | 0.0460 | <0.0001 |
| Mut2-F228L | >0.9999 | 0.0027 | >0.9999, ns |
| Mut2-del319F | <0.0001 | 0.1955, ns | <0.0001 |
